# Altered B Cell Metabolic Pathways Characterize Type 1 Diabetes Progression

**DOI:** 10.1101/2024.07.03.601778

**Authors:** Holly Conway, Dianna Perez, Mugtaba Swar-Eldahab, Jon Piganelli, Carmella Evans-Molina, Jamie Felton

## Abstract

Type 1 diabetes (T1D) results in immune-mediated destruction of insulin-producing beta cells in the pancreas. B cells have been identified as critical, pathogenic antigen presenting cells and their specificity drives disease progression. At the same time, immunosuppressive, IL-10- producing regulatory B cells (Bregs) have been shown to play protective roles in mouse models of several autoimmune diseases, including rheumatoid arthritis and multiple sclerosis. In these models, microenvironmental stimuli induce regulatory B cell differentiation. Specifically, signaling through hypoxia-inducible factor 1α (HIF-1α) drives a glycolytic flux that facilitates Breg expansion. While Breg frequencies are decreased in individuals with T1D compared to healthy controls, the role B regs play, how microenvironmental stimuli influence their differentiation, and whether this is altered in T1D progression, which is characterized by progressive, systemic hyperglycemia, are less clear. Here we examine the relationship between B cell differentiation, cellular metabolism, and HIF-1α to reveal that in a mouse model of autoimmune diabetes, B cells have distinct metabolic characteristics that change with disease progression. Further, response to hypoxia in autoimmune B cells is distinct from the response by non-autoimmune, control B cells. Together, these data suggest that dysregulated HIF signaling may skew the B cell repertoire toward inflammatory, rather than regulatory B cell subsets to drive T1D development. Consequently, HIF-1α activation to expand regulatory B cell populations may be a viable option for immune modulation.

## INTRODUCTION

Abnormalities in B cell tolerance checkpoints have been identified in individuals with type 1 diabetes (T1D), resulting in expansion of autoreactive B cells and the development of autoantibodies to islet antigens in the blood ^1–4^. In addition to autoantibody production, pathogenic roles for B cells include antigen presentation ^5–7^ and proinflammatory cytokine secretion ^8^. Emerging literature in T1D and other autoimmune diseases suggests that B cells also play protective, anti-inflammatory roles in disease progression via IL-10 production, as regulatory B cells (Bregs) ^9–12^. Clinical trials using B-cell depleting agent rituximab in individuals newly diagnosed with T1D only transiently preserved insulin secretion ^13,14^, highlighting a complex and incompletely understood role for B cells in T1D progression.

Over the last 25 years, cellular metabolism has emerged as a key modulator of immune cell function ^15–21^. Changes in tissue microenvironments prompt alterations in bioenergetic pathways, such as glycolysis and oxidative phosphorylation, which yield metabolites and biosynthetic intermediates that affect cell signaling, regulate gene expression, and polarize cell development toward pro or anti-inflammatory phenotypes ^19^. Hypoxia-inducible factors (HIFs) are key mediators of these alterations and link cellular metabolism to immune cell function ^16,22,23^.

Recently, regulation of HIF-1α, which is induced in response to microenvironment hypoxia, was shown to be critical to normal B cell development ^24^. In mouse models of experimental autoimmune encephalomyelitis and collagen-induced arthritis, HIF-1α-dependent glycolysis facilitated expansion of regulatory B cell subsets, and mice with B cell-specific deletion of *Hif1a* had exacerbated autoimmune disease ^25^. At the same time, HIF-mediated signaling has also been implicated in resolution of beta cell injury in the islet during T1D development ^26^. Roles for HIF-mediated signaling in B cell metabolism during T1D progression; however, are not known.

In the non-obese diabetic (NOD) mouse, a model of spontaneous, organ-specific autoimmunity, the development of pathogenic, autoreactive B cells has been shown to be preferentially skewed toward late transitional and marginal zone compartments, which are uniquely sensitive to environmental and homeostatic signals ^9,27,28^, suggesting their potential to be modulated by microenvironmental factors to either promote or protect against autoimmunity. The metabolic state of B cells in T1D throughout disease progression, which is marked by progressive lymphocytic infiltration of a highly vascularized islet microenvironment, in addition to progressive, systemic hyperglycemia, remains largely unknown. Understanding the links between B cell differentiation, B cell metabolism, and HIF-1α expression during T1D progression has important therapeutic implications. HIF-1α stabilizers are emerging as innovative therapy in inflammatory disorders, anemia, and wound healing ^29,30^; therefore, the development of small-molecule modulators of the HIF pathway are already well underway ^31^, which will facilitate faster transition of therapies to the clinic. Combined with the beneficial effects that have been shown on beta cell survival ^26,32^, HIF-activation is a particularly attractive target for modulation of disease development in T1D.

In this study, we seek to understand the metabolic state of B cells throughout disease progression in the NOD mouse and interrogate how HIF-1α alters B cell metabolism to impact B cell effector function. We demonstrate that metabolic programming and HIF-1α-driven responses in B cells from NOD mice are distinct from those in non-autoimmune prone mice which support HIF-1α as a potential target for tolerance induction in T1D.

## METHODS

### Mice

NOD mice were purchased from The Jackson Laboratory (Bar Harbor, ME, USA). All mice were housed and bred under specific pathogen-free conditions, and all studies were approved by the Institutional Animal Care and Use Committee of the Indiana University School of Medicine, fully accredited by the Association for Assessment and Accreditation of Laboratory Animal Care.

Both male and female mice were examined. Disease status was confirmed at the time of the experiment. Mice were considered diabetic after the first of 2 consecutive blood glucose readings > 250mg/dl.

### Cell isolation, Flow cytometry, and Abs

Freshly isolated spleens, pancreatic lymph nodes (PLNs), and pancreata were digested in 1 mg/mL collagenase P (Sigma) diluted in HBSS (Invitrogen/Life Technologies) at 37°C. Spleens and PLNs were macerated with HBSS + 10% FBS (HyClone) to inhibit collagenase activity and passed over a 70µm filter to prepare single cell suspensions. Splenic RBCs were lysed with Tris-NH4Cl. For pancreata, tissue was disrupted using an 18G needle, HBSS + 10% FBS was added to inhibit collagenase activity, and cells were passed over a 30µm filter to prepare a single cell suspension. For assessment of mitochondrial mass and polarity, cells were incubated with MitoTracker Green FM (30nM; Invitrogen) and MitoTracker Red CMXRos (100nM; Invitrogen), respectively, for 15 min at 37 °C prior to staining for extracellular markers. For glucose uptake analysis, cells were rested in glucose free cell culture media (DMEM sodium pyruvate (-) glucose (-) L-glutamine (+) [Gibco] containing 10% FBS, 1% penicillin-streptomycin, and 0.1% 2-ME; Life Technologies) for one hour prior to incubation with 60µM 2-NBDG in for 30 min at 37 °C and staining for extracellular markers using the following reagents and antibodies (BD Biosciences, Tonbo or Invitrogen): LIVE/DEAD Fixable Yellow, CD4 (RM4-5) (TONBO Biosciences), CD8 (53-6.7) (Invitrogen), CD25 (PC61.5) (TONBO Biosciences), B220 (6B2) (Invitrogen), and CD19 (1D3) (Invitrogen). Samples were acquired using an Attune NxT Flow Cytometer, and FlowJo software (Tree Star) was used for data analysis.

### B cell stimulation

Splenic B cells were purified via negative selection, using an EasySep Mouse B Cell Separation Kit (Stemcell Technologies, #19854) in accordance with manufacturer’s instructions. B cells were cultured in high glucose (4.5 g/L) DMEM + 10% FBS, 1 mM sodium pyruvate, 50 uM β- Mercaptoethanol, 2 mM L-Glutamine, penicillin (50 U/mL), streptomycin (50 ug/mL), and 10 mM HEPES at a concentration of 2 x 10^6^ cells/mL in 1 mL of medium per well in 48-well flat-bottom culture plates. 10 ug/mL LPS (Cell signaling Technology, Inc., #14011S), µ chain specific F(ab’)₂ fragment goat anti-Mouse IgM (Jackson ImmunoResearch, #115-006-075), or anti-mouse CD40 (BD Cell Analysis, #553721) was added to culture as indicated. B cells were harvested at indicated timepoints in culture and immediately lysed and homogenized in Buffer RLT plus (Qiagen).

### RT-qPCR

RNA was extracted with a Qiagen RNeasy Plus Mini Kit (Qiagen, #74136) and quantified using an Implen NP80 nanophotometer. RNA from each sample was reverse-transcribed to cDNA with a High-Capacity cDNA Reverse Transcription kit (Thermo Fisher Scientific, #74136). Real-Time PCR was performed on each sample using an Applied Biosystems QuantStudio 3 instrument with PowerUP SYBR Green Master Mix (Applied Biosystems, #A25741) and primers specific for Hif1a, Hif2a, and Actb. Primer sequences are available upon request. Ct data were exported and relative transcript levels of both Hif1a and Hif2a were calculated as described in Livak et. al^33^ after normalizing target gene Ct values to those of Actb.

### RNA Sequencing

One nanogram of total RNA per sample with RNA integrity values ≥ 7.0 was used for library preparation. SMART-Seq v4 Ultra Low Input RNA Kit for Sequencing (Takara Bio) was used for cDNA was synthesis. A dual indexed cDNA library was then prepared with use of Nextera XT DNA Library Prep Kit (Illumina). Each library was quantified, and Qubit and Agilent Bioanalyzer was used to access quality. Multiple libraries were pooled in equal molarity. The average size of the library insert was ∼300–400 base pairs (bp). The pooled libraries were denatured and neutralized before loading to the NovaSeq 6000 sequencer for 100b paired-end sequencing (Illumina). Approximately 30–40 × 106 reads per library were generated. A Phred quality score (Q score) was used to measure the quality of sequencing. More than 95% of the sequencing reads reached Q30 (99.9% base call accuracy). The generated FASTQ files were processed with the Genialis visual informatics platform (https://www.genialis.com).

### Statistical analysis

Statistical analyses were performed using GraphPad Prism 8.4.3 (GraphPad Software, La Jolla, CA). Throughout, asterisks are used to denote p values by the indicated statistical test: *p < 0.05, **p < 0.01, and ***p < 0.001.

## RESULTS

### B cell metabolism changes with type 1 diabetes progression

To understand how B cell metabolism is influenced by progressive, islet immune cell infiltration (insulitis) and systemic hyperglycemia, we evaluated metabolic signatures in B cells from NOD mice in the spleen, draining pancreatic lymph nodes (PLNs), and infiltrated islets in the pancreas, throughout T1D disease progression. In our colony, approximately 90% of female NOD mice will develop T1D by 22 weeks, and median age at diabetes development is 18 weeks for female mice; 29.5 weeks for male mice (Suppl. Fig 1). Therefore, we established three timepoints for analysis of disease progression: prior to significant islet infiltration (<8 weeks), during the window of high islet infiltration but prior to overt hyperglycemia (9-16 weeks), and with frank diabetes (defined as blood glucose >250mg/dl on two consecutive days; all mice in this group were >16 weeks). To assess glucose metabolism, glucose uptake was measured in B lymphocytes in the spleen, PLNs, and islet at each timepoint. In all tissues, glucose uptake peaked prior to diabetes onset (9-16 weeks) and decreased to levels similar to glucose uptake at < 8 weeks after development of T1D (Fig. 1). Glucose uptake was lowest in CD19+B220+ B cells, compared to CD4+ and CD8+ T subsets throughout disease progression in the spleen, PLNs, and islet (Supplemental Fig. 2), which is consistent with previous findings that resting B cells have lower energy demands and consume less glucose than resting T cells^34^. Directly ex vivo, no significant differences in glucose uptake in the spleen, compared to the PLNs or islet were identified at any point in disease progression (Fig. 1B).

**Fig. 1.**
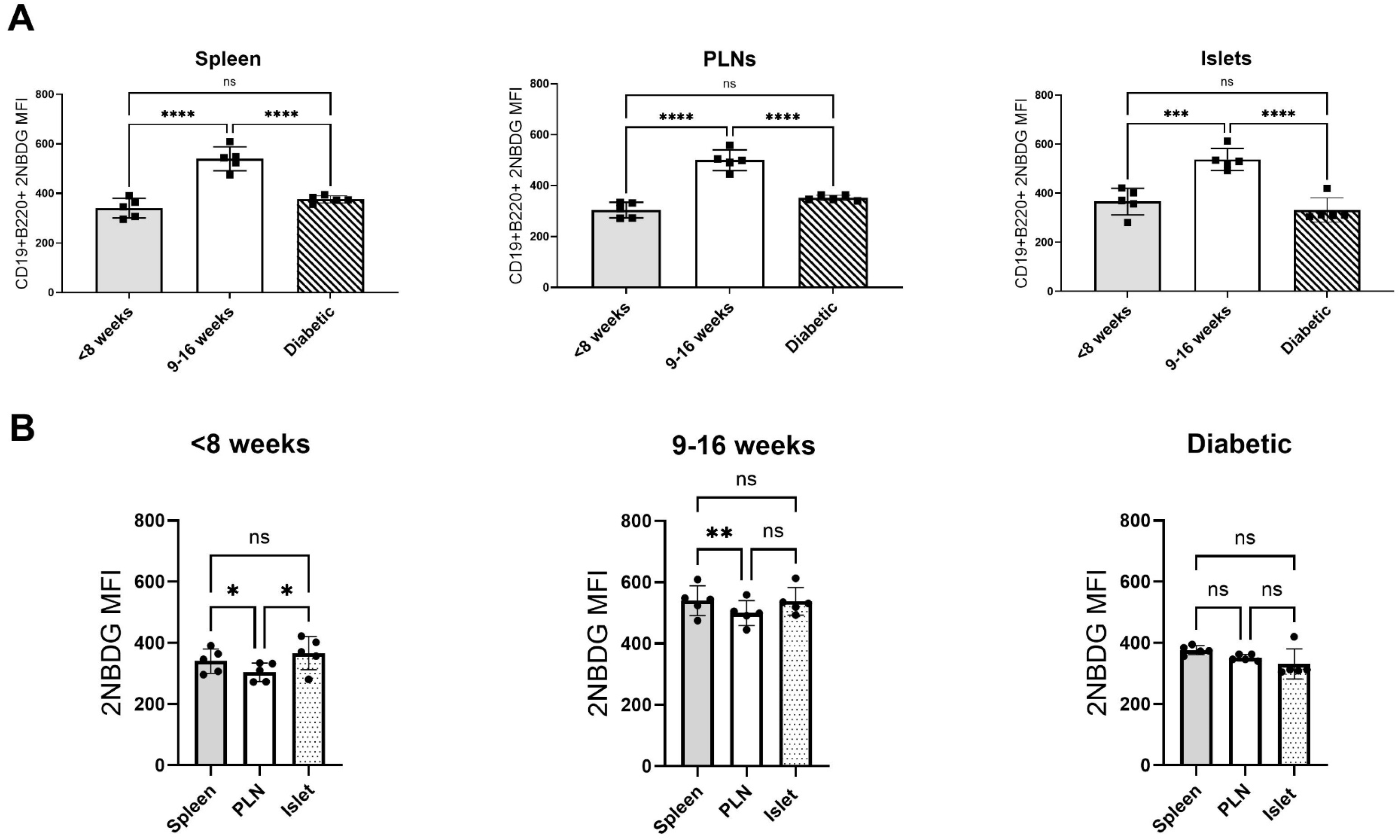
Glucose uptake is highest prior to disease onset. (A) 2NBDG was used to compare glucose uptake in CD19+B220+ B lymphocytes from the spleen, PLNs, and islets of NOD mice throughout disease progression. (B) Direct comparisons of glucose uptake in CD19+B220+ B lymphocytes in tissues from female NOD mice at different stages in disease development. *p< 0.05, **p<0.01, ***p<0.001, ****p<0.0001, one-way ANOVA with Tukey’s multiple comparison test. n= 5 for all timepoints.

As a central intermediate in oxidative energy metabolism, mitochondrial membrane potential (MMP) was measured and normalized for mitochondrial mass at the same three timepoints in B lymphocytes in the spleen, PLN, and islet throughout disease progression in NOD mice. Like patterns observed for glucose uptake, MMP increased significantly at the 9-16 week timepoint, prior to diabetes onset (Fig. 2). However, MMP remained elevated with progression to diabetes in the spleen and islets (Fig 2A) and decreased significantly in the PLNs. Tissue specificity was observed for MMP for the 9-16 week timepoint: MMP was lowest in the spleen and highest in the islet. No differences were observed between the spleen, PLN, and islets in <8 week or diabetic NOD mice (Fig. 2B). Consistent with previous literature, resting B cells had higher MMP compared to resting CD4+ and CD8+ T cells^34^ (Suppl. Fig 3A-C).

**Figure 2.**
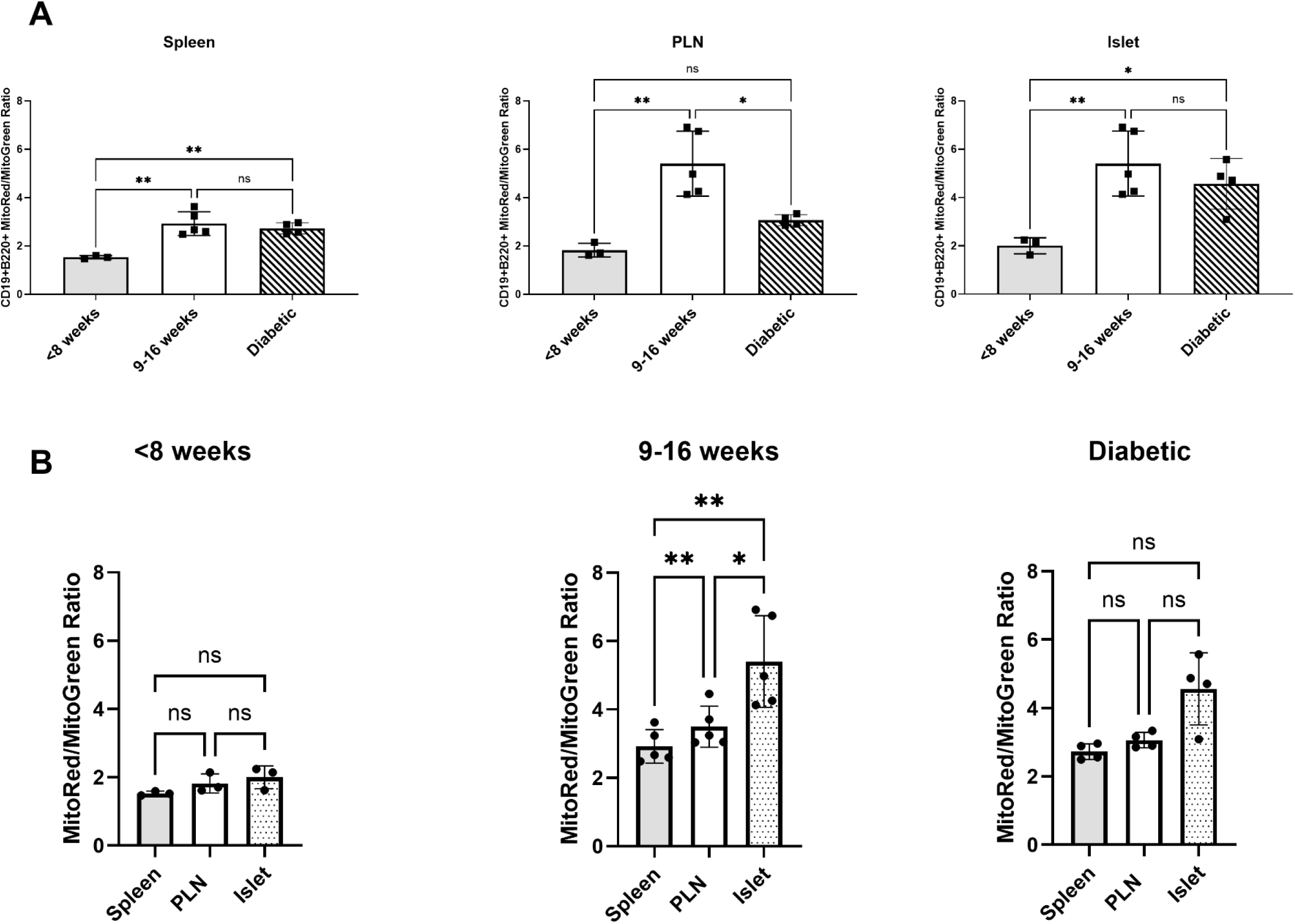
Mitochondrial membrane potential (MMP) increases most significantly in the islet prior to disease onset. (A) Mitotracker red and green were used to assess mitochondrial membrane potential and mass, respectively, in CD19+B220+ B lymphocytes from the spleen, PLNs, and islets of NOD mice throughout disease progression. (B) Direct comparison of MMP of CD19+B220+ B lymphocytes in tissues from female NOD mice at different stages in disease development. MMP normalized to mass. *p< 0.05, **p<0.01, ***p<0.001, ****p<0.0001, RM one-way ANOVA with Tukey’s multiple comparisons test.

To determine whether tissue specific changes in MMP (and absence of tissue specific changes for glucose uptake) were features of autoimmune prone lymphocytes in the NOD model, B cells from aged-matched non-autoimmune prone C57Bl6/J (B6) mice were compared to B cells from NOD mice ex vivo. Because B6 mice do have lymphocytic infiltration in the islet, we did not compare islet-infiltrating lymphocytes. Instead, we compared glucose uptake and MMP in spleen, PLNs, and non-pancreatic “other lymph nodes” (axillary, inguinal) (Fig. 3). No significant differences in glucose uptake ex vivo were observed between NOD and B6 B cells in the spleen, PLNs, or other lymph nodes (Fig. 3A). MMP, however, was significantly decreased in non-pancreatic lymph nodes in NOD mice compared to B6 mice (Fig. 3B). Tissue specific changes were not observed for glucose uptake in B6 B cells in the spleen, PLNs, or other lymph nodes. Unlike NOD B cells, differences between MMP in the spleen and PLN were not significantly different. However, MMP was significantly higher in B6 B cells from other lymph nodes, compared to B6 B cells from the spleen and PLN (Fig. 3C, D). Given peak metabolic activity identified during the period that preceded disease onset, all mice used in subsequent experiments were 9-16 weeks old.

**Figure 3.**
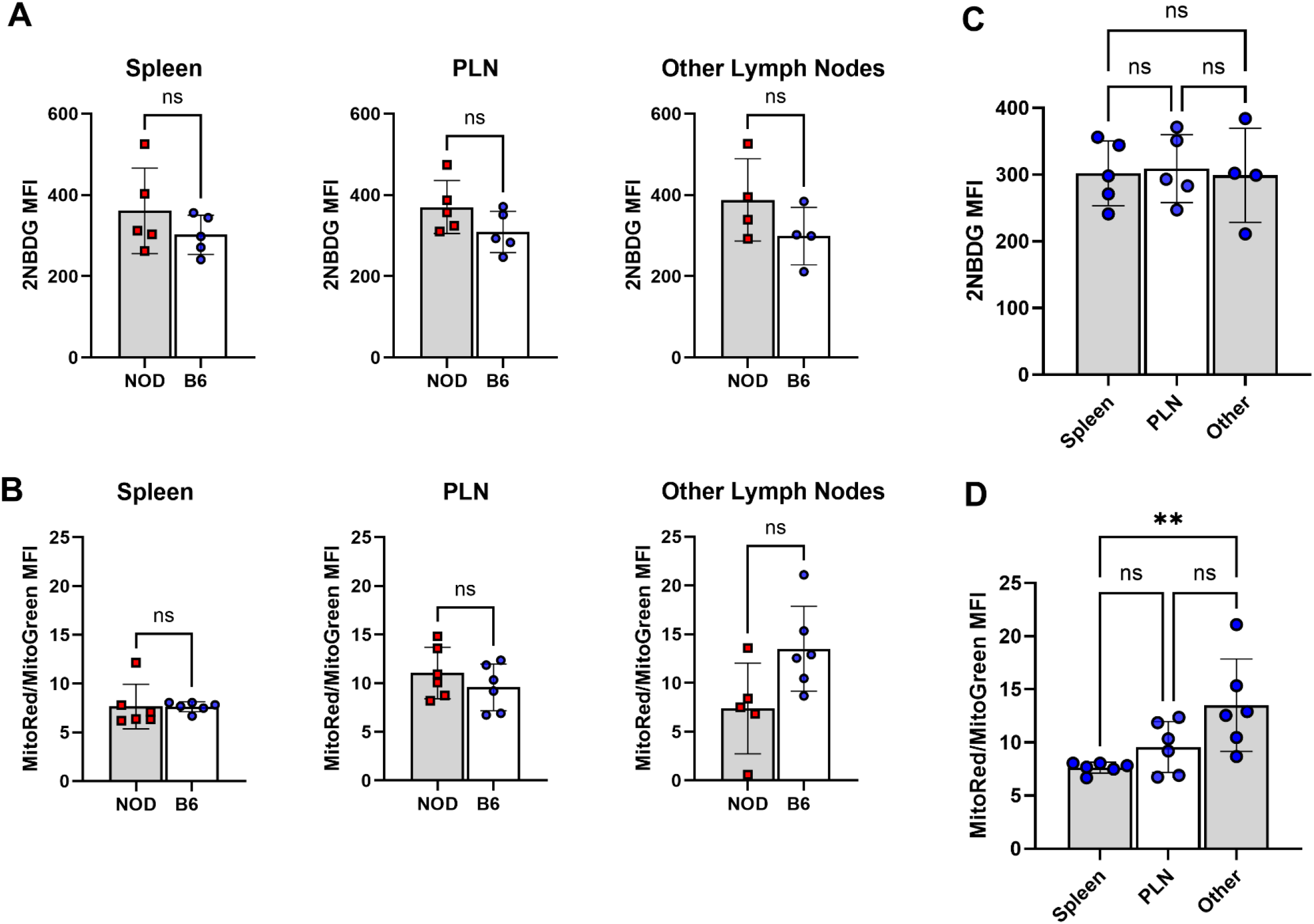
Glucose uptake and mitochondrial membrane potential (MMP) are not different between NOD and B6 B cells ex vivo. (A) 2NBDG was used to compare glucose uptake in B cells from the spleen, PLNs, and non-pancreatic "other lymph nodes" in 14-16 week NOD and B6 mice. (B) MitoTracker red and green were used to assess MMP and mass, respectively, in B cells from the spleen, PLNs, and non-pancreatic "other lymph nodes" 14-16 week NOD and B6 mice. *p< 0.05, **p<0.01, ***p<0.001, ****p<0.0001, two-tailed unpaired t-test. (C) Glucose uptake in B cells from the spleen, PLN, and non-pancreatic "other lymph nodes" were compared in B6 mice. (D) MitoTracker red and green were used to assess mitochondrial membrane potential and mass, respectively, in B cells from the spleen, PLNs, and non-pancreatic "other lymph nodes in B6 mice. MMP normalized to mass. Each point represents an individual mouse. *p< 0.05, **p<0.01, ***p<0.001, ****p<0.0001, One-way ANOVA with Tukey’s multiple comparison test.

### Activation reveals distinct metabolic phenotypes in autoreactive B cells

Because activation has been shown to drive changes in cellular metabolic profiles, we next determined whether differences between NOD and B6 B cells could be detected after activation. Splenic B cells were isolated from spleens of 14-16 week NOD and B6 mice and activated for 72 hours with 10 µg/mL of anti-CD40. While there were no differences at baseline, when stimulated, glucose uptake was significantly higher in NOD compared to B6 B cells (Fig. 4A).

**Figure 4.**
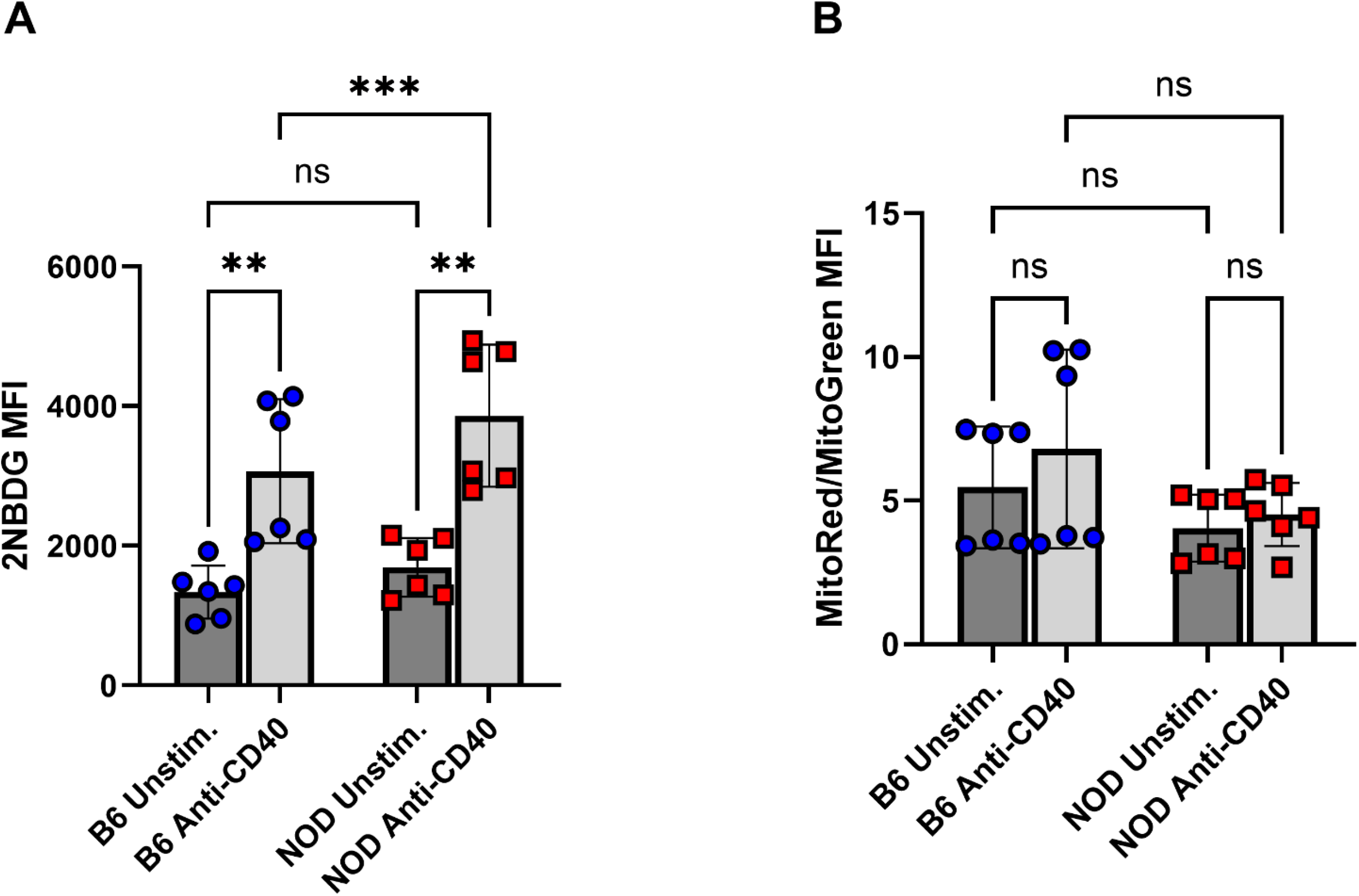
Glucose uptake is increased in activated NOD B cells. Splenic B cells were isolated from spleens of B6 and NOD mice and activated for 72 hours with anti-CD40. Glucose uptake was measured using 2NBDG (A). Mitochondrial membrane potential (MMP) was measured using MitoTrackers red and green to assess MMP and mass, respectively (B). MMP was normalized to mitochondrial mass. Each point represents an individual mouse. *p<0.05, **p<0.01, ***p<0.001, ****p<0.0001, one-way ANOVA with Tukey’s test for multiple comparisons.

Conversely, no significant changes were detected in MMP potential at baseline or after stimulation with anti-CD40 in either B6 or NOD B cells (Fig. 4B). Together, these data suggest a unique metabolic phenotype in NOD B cells characterized by oxidative phosphorylation ex vivo throughout all stages of disease progression, and exaggerated glucose uptake in response to B cell receptor stimulation.

### Hypoxia-mediated responses are divergent in autoimmune B cells

As a major transcription factor that governs cellular metabolic responses to changes in the microenvironment, HIF-mediated adaptive responses have been shown to be critical to beta cell recovery after insult^26^ as well as shaping the bioenergetics of immune cell responses^35^. HIF-1α plays critical roles in physiologic B cell development^24^, specifically, in the development of regulatory B cell subsets. HIF-1α- mediated glycolytic flux has been shown to drive regulatory B cell expansion in collagen-induced arthritis (CIA) and experimental autoimmune encephalomyelitis (EAE), and HIF deficiency led to exacerbated autoimmunity^25^. Regulatory B cell responses have been shown to be impaired in human T1D and NOD mouse models^36^ ; therefore, we asked whether dysregulated HIF signaling may impair regulatory B cell responses in NOD mice. B cells from NOD and B6 mice were cultured in room air or under hypoxic conditions to drive HIF-1α stabilization, and transcriptomes in NOD and B6 B cells were compared. In CIA and EAE, regulatory B cell development required a HIF-1α driven glycolytic flux, and HIF deficiency reduced expression of HIF-1α-targeted glycolytic genes *Glut1, Pkm2, Hk2, Ldha, Pdk1, and Gpi1,* impaired regulatory B cell expansion, and consequently, exacerbated autoimmune disease^25^. In NOD B cells cultured in hypoxic conditions, HIF-1α- targeted glycolytic mRNA expression was reduced compared to non-autoimmune B6 B cells (Fig. 5A). To investigate whether reduction in HIF-driven glycolytic gene expression had implications for regulatory B cell development in NOD mice, mRNA expression of 19 common marker genes for regulatory B cells^37^ were compared from B cells cultured in room air and in hypoxia. Of these 19 identified genes, most were downregulated in hypoxic NOD B cells (Fig. 5B). Interestingly, differential expression analysis of transcripts from NOD B cells cultured in room air or hypoxic conditions demonstrated significantly decreased expression of *Cd274*, which encodes PD-L1, in response to hypoxia (Fig. 5C). PD-L1 has been shown to be highly expressed on some subsets of regulatory B cells^38^. In addition, four of the 24 genes that comprise the hallmark IL6/JAK/STAT3 signaling Molecular Signatures Database (MSigDB) gene set^39^ are also downregulated in hypoxic NOD B cells. In B6 B cells, HIF-1α and STAT3 have been shown to cooperatively regulate *Il10* transcription^25^; therefore, together, these data suggest that the cellular response to hypoxia is dysregulated and/or insufficient in B cells from NOD mice.

**Figure 5.**
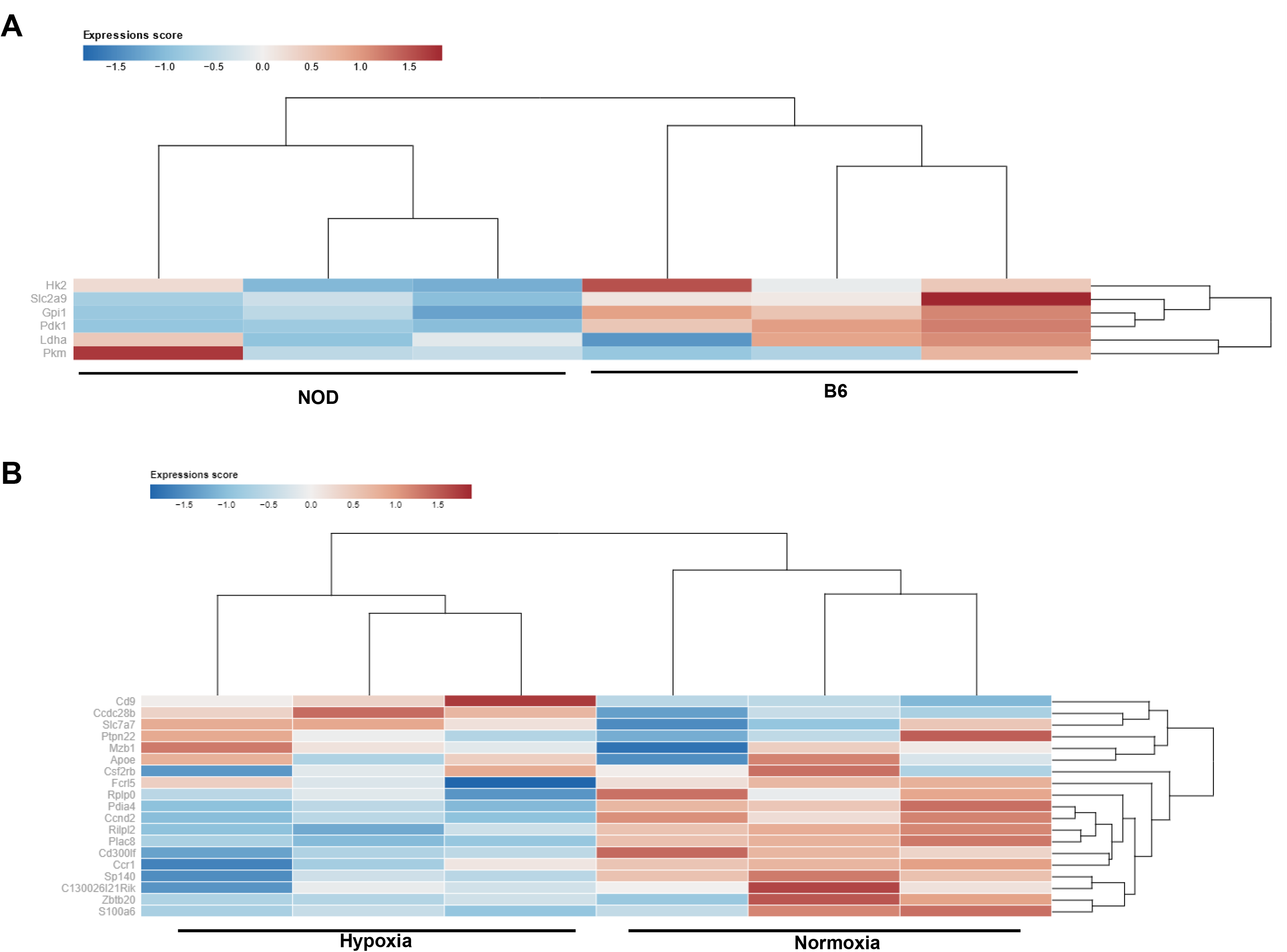
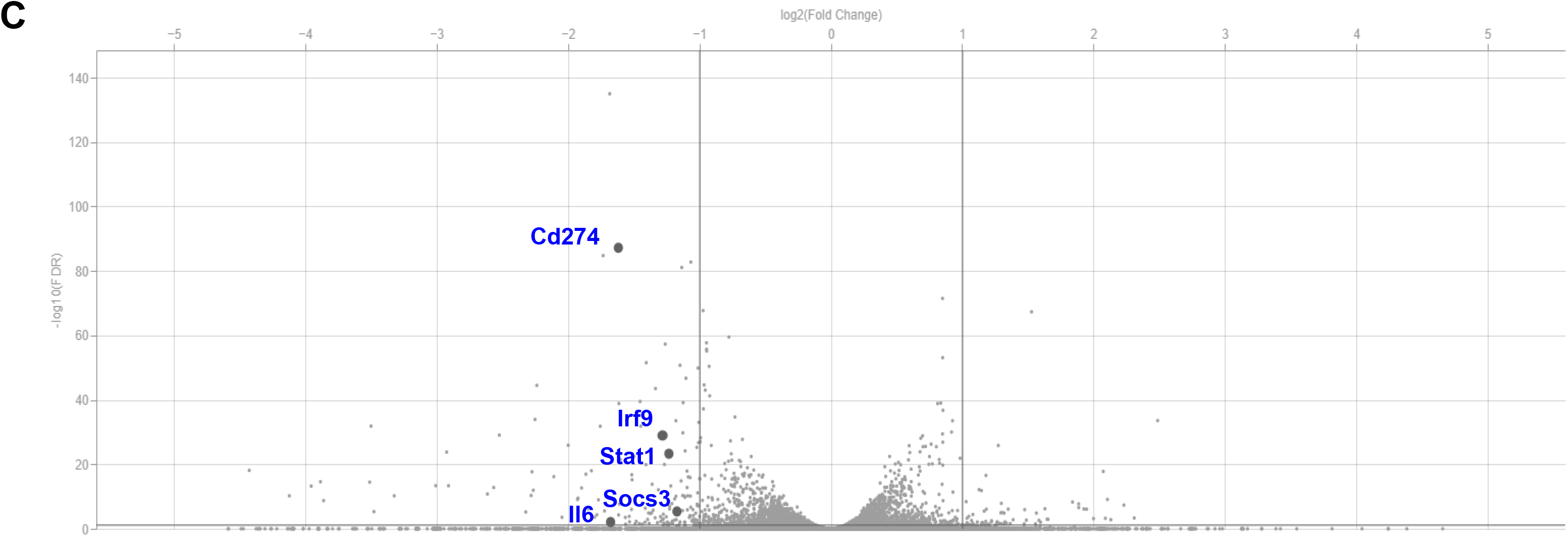
Distinct glycolytic and regulatory B cell transcripts characterize NOD and B6 response to hypoxia. (A) Heatmap showing row-wise scaled expression using Z-score based on the raw gene expression of selected glycolytic genes for B cells from NOD and B6 mice (n=3 for each group, 12-16 week female mice) cultured for 24 hours at 5% pO2. Expression type prior to scaling is TPM. The color scale goes from -1.38 to 1.84. (B) Heatmap showing row-wise scaled expression using Z-score based on the raw gene expression of selected regulatory genes for B cells from NOD mice cultured for 24 hours at 5% pO2 or room air (n=3 for each group, 12-16 week female mice). Expression type prior to scaling is TPM. The color scale goes from -1.99 to 1.81. (C) Differential expression of in NOD B cells cultured in hypoxia or room air for 24 hours. X-axis indications the log2 value of fold change in differential expression. The FDR on the y-axis is the probability that the gene is differentially expressed.

Indeed, follow-up evaluation of *Hif1a* mRNA expression also suggested insufficient HIF induction in the NOD compared to B6 B cells. In response to stimulation with B cell mitogens (LPS and anti-IgM), B cells from B6 mice have been shown to increase Hif*1a* mRNA expression^25^. To determine whether *Hif1a* mRNA expression is similarly induced in NOD B cells, we stimulated splenic NOD B cells with LPS (10ug/ml), anti-IgM (10ug/ml) or anti-CD40 (10ug/ml) and compared mRNA expression to aged-matched B6 mice (Fig. 6). Compared to B6 B cells, *Hif1a* expression in NOD B cells was lower at all time points for all stimuli. *Hif1a* expression was significantly decreased in response to anti-IgM at 24 hours (Fig, 6B). *Hif1a* was also significantly decreased in response to anti-CD40 at 8 and 24 hours (Fig. 6C).

**Fig. 6.**
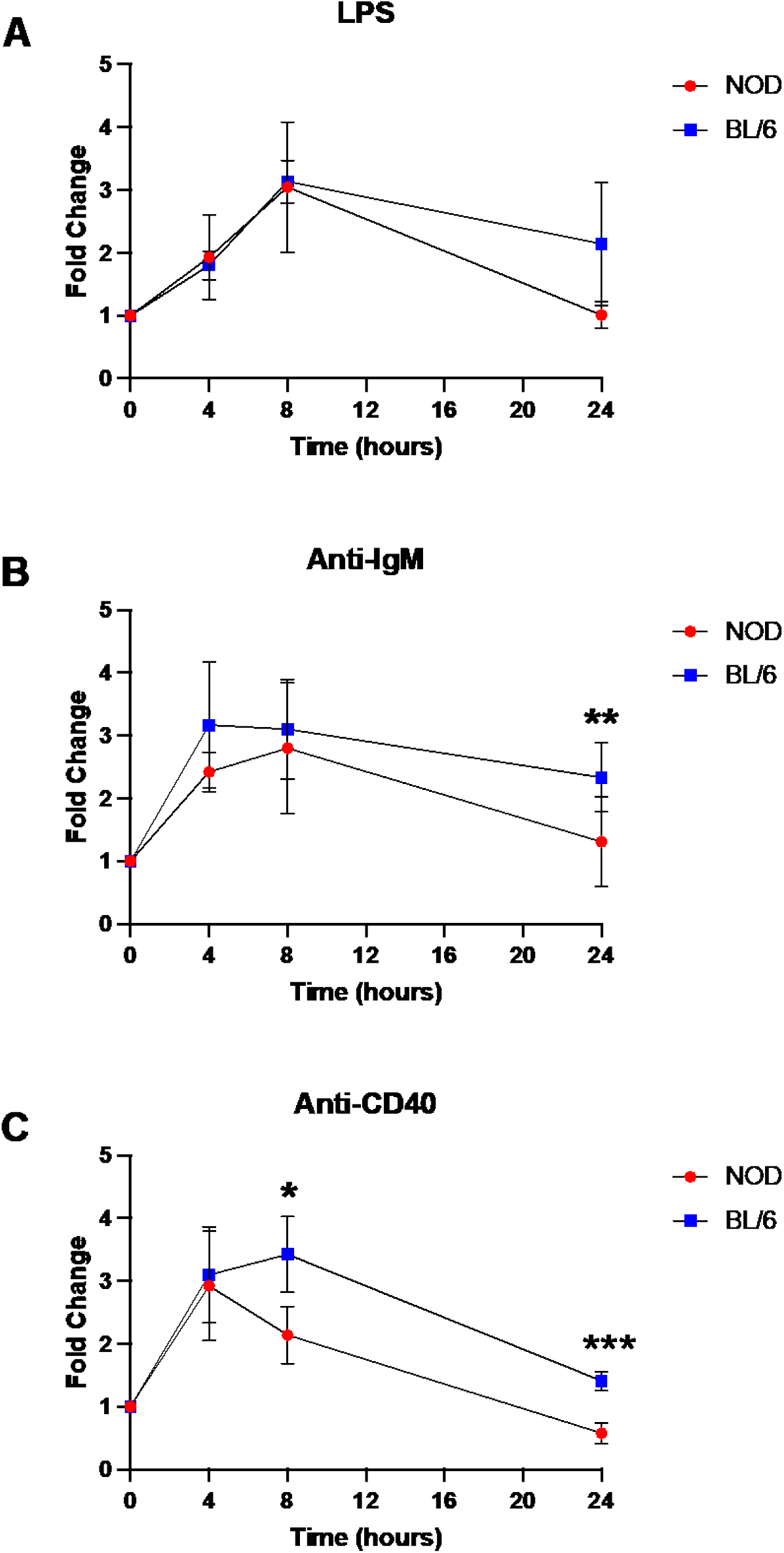
HIF-1⍺ expression is altered in activated NOD B cells. Splenic B cells were isolated from spleens of 12-16 week female B6 and NOD mice (n=4 mice per strain). B cells were activated with (A) LPS, (B) Anti-IgM, and (C) Anti-CD40 and gene expression was measured at the indicated time points. *p< 0.05, **p<0.01, ***p<0.001, ****p<0.0001, multiple, paired t-test.

## DISCUSSION

Immune modulation for type 1 diabetes prevention has been a goal since the discovery that T1D was immune mediated in the early 1980s. The role that cellular metabolism plays in altering immune cell function has become increasingly recognized, expanding the armamentarium of therapeutic targets for disease modulation. B cells have been shown to play critical roles in T1D development as antigen-presenting cells, antibody producing cells, and more recently, suppressive, regulatory B cells, highlighting multiple fates for B cells during development and multiple opportunities to modulate immune responses^6,36,40–44^. However, only B cell depletion has been tested in clinical trials to alter T1D disease course^13,14^. Subsets of B cells with regulatory properties have been shown to be protective in mouse models of autoimmune disease^45–48^. Alterations in B cell subsets have been identified in the blood of individuals with autoimmune diabetes^10,49^, suggesting that in both human and mouse models, B cells play a role an important role in T1D pathogenesis. Therefore, targeting specific populations of autoreactive B cells for depletion, or regulatory B cells for expansion, represent promising alternatives to systemic depletion therapies. The unique and distinct metabolic features of autoimmune B cells from NOD mice expand the armamentarium of agents for selective, therapeutic targeting.

Therefore, understanding B cell function during disease progression in T1D is an understudied opportunity to develop novel and effective interventions.

Compelling evidence from other autoimmune disease models suggests that B cell response to hypoxia has important implications for development of regulatory B cells^25,50,51^. In T1D, B cells are required for disease development, and autoimmune B cells have been shown to accumulate in compartments exquisitely sensitive to environmental changes^9^. While cellular metabolism has been shown to influence B cell development in other models of disease, B cell cellular metabolism during T1D progression has not been well studied. Here, we demonstrate three important concepts that highlight new opportunities for targeted immune modulation in T1D: 1) B cell metabolism changes with T1D progression, 2) B cell metabolic responses in autoimmune B cells are distinct from non-autoimmune B cells, and 3) alterations in both HIF expression and HIF-mediated responses are observed in NOD compared to B6 B cells with implications for regulatory B cell development.

It is not surprising that cellular metabolism changes with disease progression. In T1D, gradual chronic destruction of insulin-producing beta cells in the islet leads to lymphocytic infiltration (insulitis), leading to islet inflammation. At the same time, the development of progressive hyperglycemia alters systemic metabolism; both of which have implications for immune cell function^52,53^. Understanding which pathways are altered, and at which timepoints, is a prerequisite for targeting metabolic processes in immune cells to alter their function, and the timing of strategies to target cellular metabolism are likely to be as important as the targets themselves. Simultaneous increases in both glucose uptake and mitochondrial polarity in NOD B cells suggest that glycolysis and oxidative phosphorylation are both maximal during periods of islet inflammation, immediately prior to diagnosis and then decrease following T1D onset.

Dimethyl fumarate is an anti-inflammatory agent that inhibits the glycolytic enzyme glyceraldehyde 3-phosphate dehydrogenase (GAPDH)^54^, and has been shown to preferentially inhibit pro-inflammatory cytokine-producing B cells in multiple sclerosis^55^. This may be a viable agent to delay onset of T1D prior to diagnosis, but if initiated after disease onset, when glucose uptake decreases, may show little benefit. Similarly, metformin is an agent that, in addition to its effects on insulin sensitivity, has been shown to also serve as an anti-inflammatory agent by altering immune cell metabolism, primarily through inhibition of the mitochondrial electron transport chain^56^. While clinical studies that have investigated the effects of metformin for adjunctive treatment in T1D have been largely equivocal and have failed to show robust improvement in metabolic control^57–59^. Many of these, however, have focused on metformin as an insulin sensitizer after diagnosis or in individuals with significant insulin resistance. Given its anti-inflammatory effects driven by targeting oxidative phosphorylation in immune cells, it is possible that when tested when oxidative phosphorylation is most active, metformin could play a role in delaying ongoing beta cell destruction. These studies begin to paint a picture of evolving cellular metabolic phenotypes in immune cells in T1D that, when more clearly understood, have the potential to repurpose older agents for immune modulation when used at different times during disease progression.

Selective targeting of inflammatory, self-reactive effector cells remains a challenge in immune modulation of T1D. While the anti-CD3 agent teplizumab has had success in delaying T1D onset^60,61^, the risks associated with systemic targeting of the T cell population limit its use and raise safety concerns regarding repeated dosing. Antigen-specific therapy, the antidote to global targeting of cell populations, remains a lofty and unrealized goal, in part due to the diversity and expansion of islet antigens with disease progression. Here, by demonstrating unique metabolic phenotypes in autoimmune-prone NOD B cells compared to non-autoimmune prone B6 B cells, we suggest that immunometabolism-specific therapy is an alternative approach to selective targeting of self-reactive effector cells in T1D. The exaggerated increase in glucose uptake observed with activation in NOD B cells compared to B6 B cells suggests potential to modulate glucose metabolism in order to suppress inflammatory responses, similar to what has been done in oncology^62^. The next step, therefore, is specific analysis of key enzymes or pathways that predominate autoimmune B cell metabolism for identification of viable therapeutic targets.

Benefits to further investigation of glucose metabolism in B cells is that a handful of anti-cancer drugs that target glucose metabolism have already been developed, which, if can be repurposed for autoimmune indications, will facilitate faster translation from bench to bedside.

As role for regulatory B cells in T1D development is increasingly recognized, so too is the possibility of expansion of regulatory cells for immune modulation, in addition to repression of inflammatory cells. The HIF pathway is recognized for its role in controlling cellular metabolism gene expression, thereby controlling a cell’s function^63^. The identified role for HIF-1α in regulatory B cell expansion in other models of autoimmunity prompted our investigation of a role for HIF-1α during T1D development. Importantly, we have previously shown that autoreactive, anti-insulin B cells have enhanced IL-10 production compared to polyclonal B cells, suggesting the capacity for polarization into regulatory rather than inflammatory subsets with the right stimuli or environmental triggers^9^. Indeed, we found evidence that the HIF-dependent glycolytic flux that drives regulatory B cell expansion in models on the B6 background is likely dysregulated in NOD mice. HIF induction failed to upregulate key genes in the glycolytic pathway. At the same time, the majority of common regulatory B cell marker genes were downregulated in NOD B cell in response to hypoxia. With stimulation, HIF expression increases less in the NOD compared to B6 B cells, pointing to insufficient HIF induction as a liability to regulatory B cell expansion in the NOD mouse. If this is the case, then targeting HIF for activation and/or modulating how NOD B cells respond to HIF induction by targeting key elements in its signaling pathway represent an viable option for regulatory B cell expansion.

This is supported by examples of pharmacological activation of HIF using prolyl hydroxylase domain-containing protein (PHD) inhibitors to control inflammation in renal anemia and other inflammatory disorders, including roxadustat and daprodustat^63^. These successes will ultimately expedite translation of their use in other disease models. At the same time, additional beneficial roles have been identified for HIF activation in the management of diabetes. There is evidence of HIF-1α repression in STZ-models of diabetes that worsens prognosis of fungal infections (precedent for repressed HIF), in addition to evidence for HIF-1α repression secondary to hyperglycemia^64,65^. HIF-1α has been shown to promote autophagy and protect beta cell survival in the setting of inflammatory stress^32^. Therefore, HIF activation has potential for anti-inflammatory through driving regulatory B cell expansion, we well as support for beta cell health.

Our study has several limitations. We acknowledge that it is primarily descriptive. Further mechanistic analysis into key enzymes and pathways that drive the differences observed in autoimmune and non-autoimmune B cells are imperative for understanding how modulate glucose metabolism for therapeutic efficacy. Now that differences in glucose uptake have been identified, analysis of what happens to glucose in the setting of increased uptake without changes in oxidative phosphorylation. Real-time bioenergetic analysis will be key in unraveling these questions.

In summary, we provide evidence for distinct metabolic pathways that are active in autoimmune B cells in the NOD mouse, which support selective targeting of cellular metabolism to control immune cell function and change disease course in T1D. Based on previous data and our demonstration of dysregulated responses to HIF induction in the NOD mouse, HIF activation appears to be a reasonable place to start. When implemented at the right time during disease progression, regulation of HIF-1α in B cells has the potential to change the trajectory of disease in individuals at risk.

