## Supplementary Figures for "Altered B Cell Metabolic Pathways Characterize Type 1 Diabetes Progression"

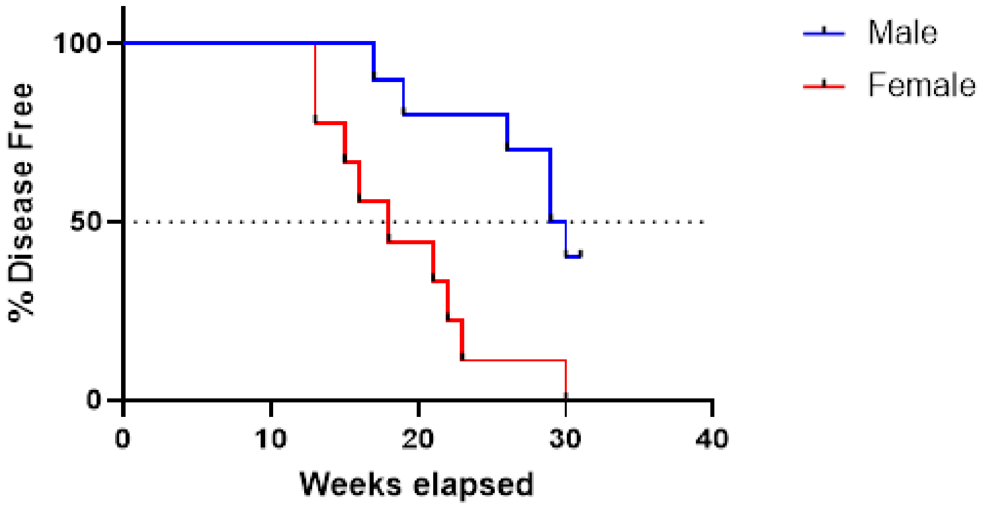


**Supplementary Figure 1. Diabetes incidence in our NOD colony.** Female (red) and male (blue) NOD mice were monitored weekly for diabetes progression. Median survival was 18 weeks and 29.5 weeks for female and male mice, respectively. n = 10 male mice, n= 10 female mice.

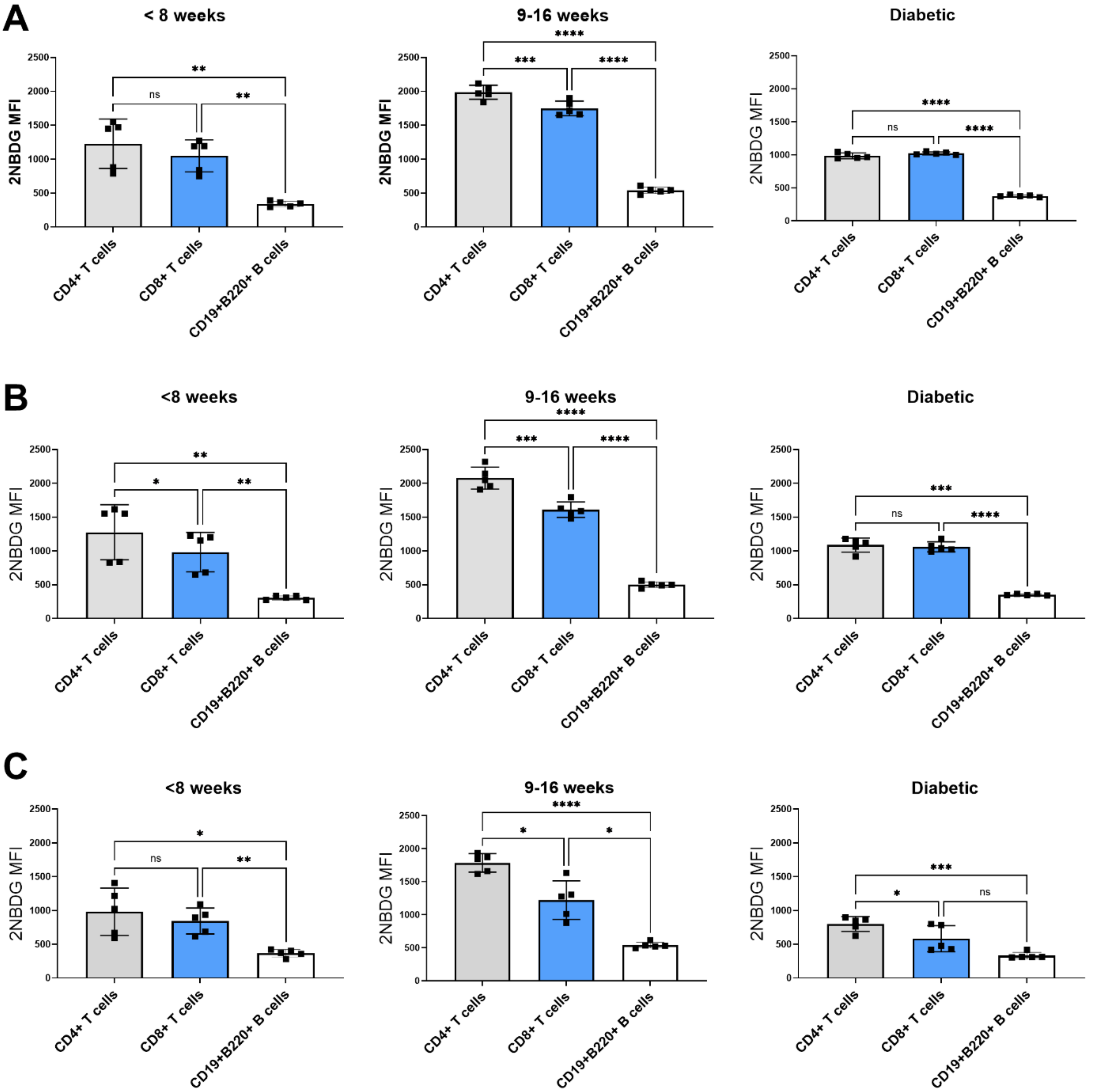


**Supplementary Figure 2. Glucose uptake is lowest in B cells throughout disease progression.** 2NBDG was used to compare glucose uptake in lymphocyte subsets from the spleen (A), pancreatic lymph nodes (B) and islets (C) of NOD mice throughout disease progression. *p< 0.05, **p<0.01, ***p<0.001, ****p<0.0001, Rm one-way ANOVA with Tukey's multiple comparisons test.

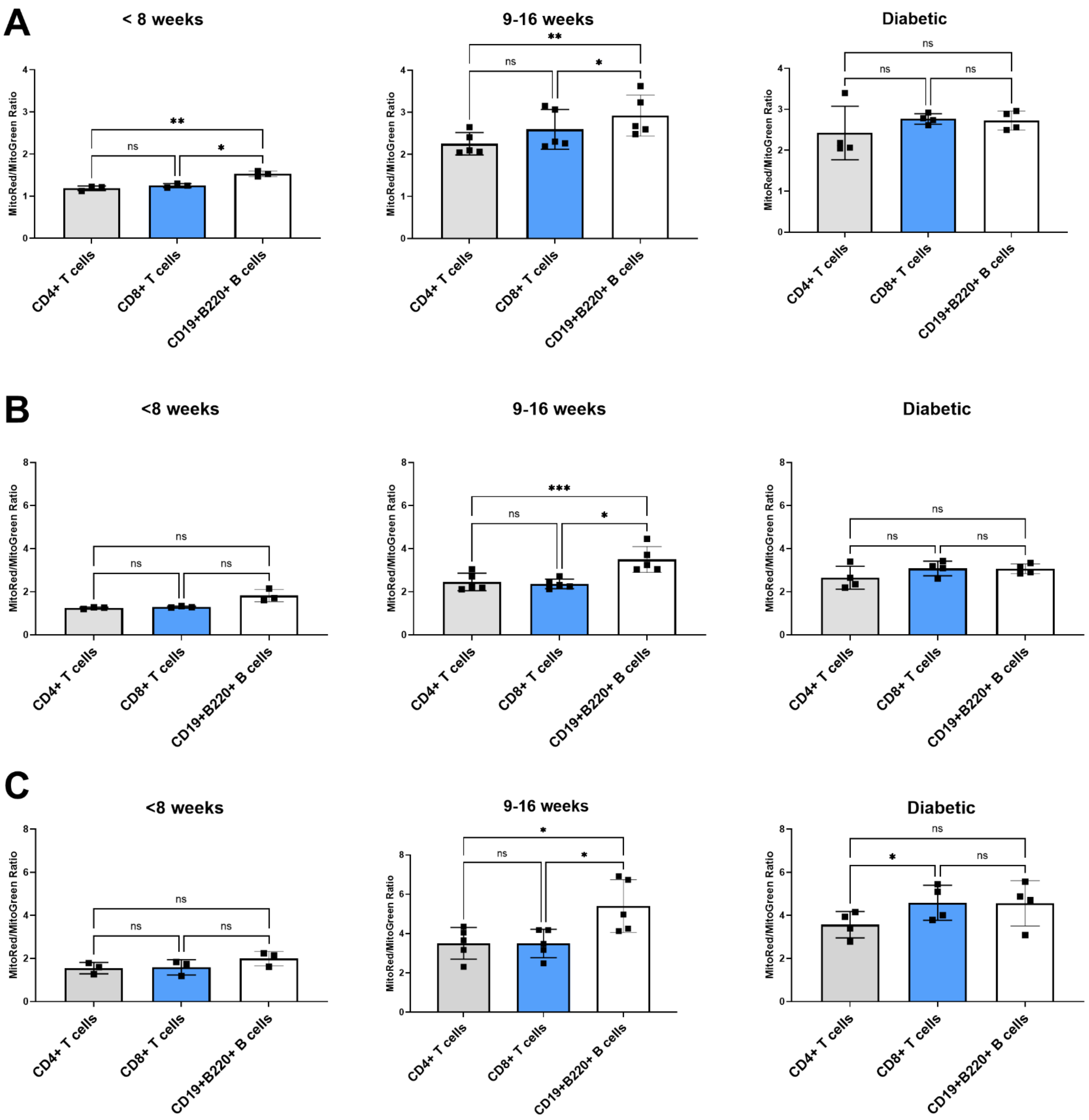


**Supplementary Figure 3. MMP is significantly higher in B cells prior to disease onset.** Mitotracker red and green were used to assess mitochondrial membrane potential and mass, respectively, in lymphocyte subsets from the spleen (A), pancreatic lymph nodes (B) and pancreas (C) of NOD mice throughout disease progression. Changes in MMP normalized to mass. *$p < 0.05$, **$p<0.01$, ***$p<0.001$, ****$p<0.0001$, RM one-way ANOVA with Tukey's multiple comparisons test.
